# Modified antibiotic adjuvant ratios can slow and steer the evolution of resistance: co-amoxiclav as a case study

**DOI:** 10.1101/217711

**Authors:** Richard C. Allen, Sam P. Brown

## Abstract

As the spread of antibiotic resistance outstrips the introduction of new antibiotics, reusing existing antibiotics is increasingly important. One promising method is to combine antibiotics with synergistically acting adjuvants that inhibit resistance mechanisms, allowing drug killing. Here we use co-amoxiclav (a commonly used and clinically important drug combination of the β-lactam antibiotic amoxicillin and the β-lactamase inhibitor clavulanate) to ask whether treatment efficacy and resistance evolution can be decoupled via component dosing modifications.

A simple mathematical model predicts that different ratios of these two drug components can produce distinct evolutionary responses despite similar initial levels of control. We test this hypothesis by selecting *Escherichia coli* with a plasmid encoded β-lactamase (ESBL CTX-M-14), against different proportions of amoxicillin and clavulanate. Consistent with our theory, we found that while resistance evolved under all conditions, the component ratio influenced both the rate and mechanism of resistance evolution. Specifically, we found that the current clinical practice of high amoxicillin to clavulanate ratios resulted in the most rapid failure due to the evolution of gene dosing responses. Increased plasmid copy number allowed *E. coli* to increase β-lactamase dosing and effectively titrate out the low quantities of clavulanate, restoring amoxicillin resistance. In contrast, we found high clavulanate ratios were more robust - plasmid copy number did not increase, although porin or efflux resistance mechanisms were found, as in all drug ratios. Our results indicate that by changing the ratio of adjuvant to antibiotic we can slow and steer the path of resistance evolution. We therefore suggest the use of increased clavulanate dosing regimens to slow the rate of resistance evolution.

## Introduction

The current crisis of antibiotic resistance is grounded in the ability of bacterial pathogens to rapidly evolve and adapt to novel stressors like antibiotics (1). Even for the same drug many different mechanisms confer resistance, often with varying transmissibility, costs and cross resistances (2-5). Examples of resistance have been found for all currently used antibiotics and recently clinicians have begun to face pathogens that are resistant to all available antibiotics (4, 6-8).

In addition to the ongoing search for new drugs (9), an important direction in combating resistance is the restoration of antibiotic sensitivity to existing drugs via the use of anti-resistance compounds or adjuvants (10-12). Antibiotic adjuvants are compounds that do not affect the growth of bacteria on their own but instead enhance the activity of antibiotics, by inhibiting mechanisms of resistance (13, 14). For example β-lactamase inhibitors prevent β-lactamase enzymes from degrading β-lactam antibiotics (15). β-lactamse mediated resistance is especially problematic for gram negative pathogens where these enzymes are common and disseminated on plasmids (10, 16, 17). Therefore β-lactamase β-lactamase inhibitor (BLBLI) combinations will restore β-lactam sensitivity of β-lactamse carrying strains without using novel antibiotics or antibiotics of last resort like carbapenems (16).

In this study we use co-amoxiclav (brand name Augmentin), a BLBLI combination of amoxicillin and clavulanate (clavulanic acid) that has been widely used globally since 1981 (18), and is on the WHO list of essential medicines (19). Amoxicillin is a bacteriocidal β-lactam antibiotic that inhibits synthesis of the bacterial cell wall (17). The adjuvant clavulanate has a similar structure to β-lactam antibiotics and thus acts as a competitive inhibitor of many β-lactamase enzymes (15). By preventing amoxicillin cleavage, clavulanate suppresses the resistance phenotype making amoxicillin effective again.

Despite the efficacy of BLBLI combinations like co-amoxiclav resistance is still possible, either by altering the effect of amoxicillin or the effect of clavulanate. Clavulanate is ineffective against resistance mechanisms that don’t involve β-lactamase expression. Thus direct resistance to amoxicillin, via altered penicillin binding protein structure, reduced porin expression or increased efflux pump expression can lead to resistance to co-amoxiclav (15). On the other hand increased production of lactamase enzymes overwhelm the clavulanate (20) and inhibitor resistant β-lactamase enzymes reduce (or abolish) the effect of clavulanate (21).

Despite the recent interest in adjuvants, the relative doses of the components in adjuvant therapies have received little attention, with clinical amoxicillin: clavulanate ratios varying from 2:1 to 16:1 (22), with an increase in amoxicillin more recently to combat resistance (18). Here we mathematically model and empirically map the synergy between amoxicillin and clavulanate in controlling a population of β-lactamase expressing *E. coli*. We then go on to demonstrate that drug ratios that are equally effective in their initial levels of control can produce distinct evolutionary responses. Specifically, we find that current high amoxicillin ratios lead to the rapid evolution of resistance via increased β-lactamase expression, while low amoxicillin ratios with equal initial efficacy are more robust, and maintain the efficacy of our meagre pool of β-lactamase inhibitors. We therefore suggest the use of increased clavulanate dosing regimens to slow the rate of resistance evolution.

## Results

### Theoretical model

We begin by developing a simple qualitative model for bacterial population dynamics under the control of co-amoxiclav as examples of β-lactam antibiotics and a β-lactamase inhibiting adjuvants (Figure S3, Text S1). The model predicts that the two compounds will show strong synergy when controlling a pathogen with an existing β-lactamase resistance gene (Figure 1a), as is seen in our experimental results (Figure 1b). We next introduce two resistance mechanisms to the model, direct (non-β-lactamase) mediated resistance via target/permeability mutations (fig 2a), or β-lactamase over-production (fig 2b) and ask how co-amoxiclav component dosing regimens impact selection for each mechanism.

**Figure 1:**
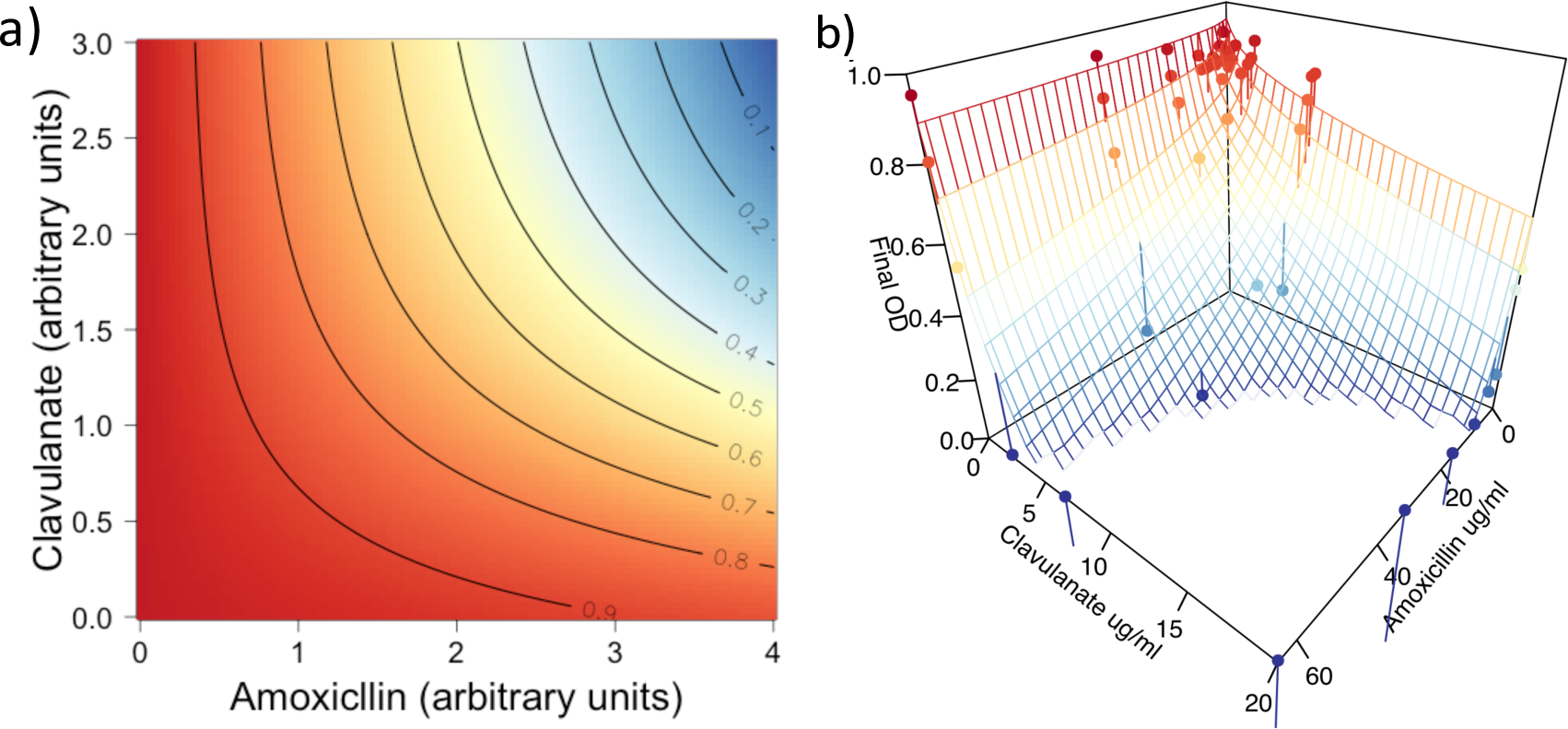
A simple model of amoxicillin and clavulanate action captures observed synergy. a) We model the dynamics of bacterial density *B* under the influence of amoxicillin *A* and clavulanate *C* as 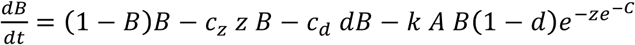 (for details, see Text S1). Carrying capacity of the system is normalised to one, so equilibrium density will be one in the absence of drugs and costly resistance. Parameters capture resistance to amoxicillin via β-lactamase over-production (*z=*2.3). Other parameters are *k*=0.3, *d*=0.1, *c*_*d*_=0.05 and *c*_*z*_=0.005. b) This model approximates the synergistic inhibition of growth seen in *E. coli* expressing a β-lactamase. The surface is the prediction of the fitted linear model (Figure S1).

**Figure 2:**
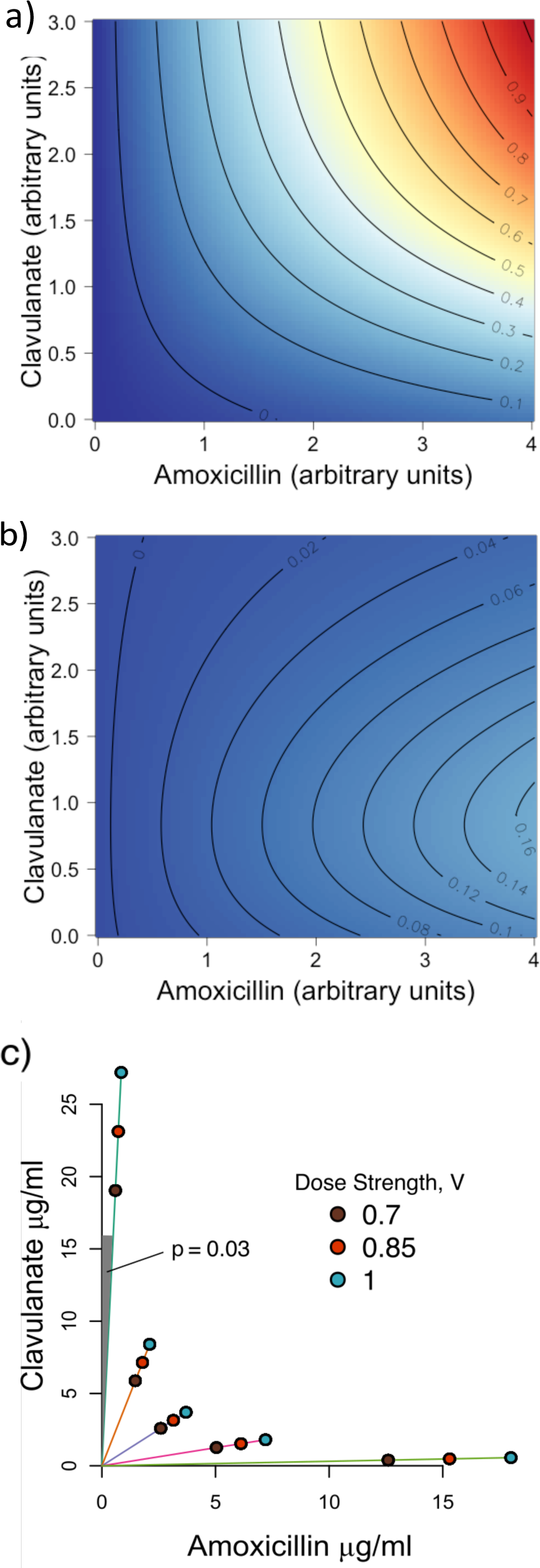
Drug ratios steer adaptation towards different resistance mechanisms in a simple model. a) The growth benefit 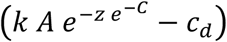 of direct (*d*, not mediated by β-lactamase) resistance to amoxicillin is directly proportional to the inhibition of bacterial growth in the model (Figure 1a). b) The growth benefit 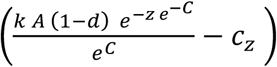 of increased β-lactamase production (*z*) can be greatly reduced without affecting equilibrium density or benefit of direct resistance by reducing the proportion of amoxicillin. Parameter values are the same as in Figure 1a. See Text S1 for derivation of equations. c) We chose 5 drug ratios (different coloured lines from the origin) which according to our model and data (Figure 1) should impose similar inhibition (at 3 different strengths of inhibition) but select for different resistance mechanisms.

The model predicts that non-β-lactamase resistance mutations will be selected in proportion to the efficacy of the combination treatment (Figure 2a). In contrast, β-lactamase over-production mutants show a more interesting pattern with maximal selection biased towards high amoxicillin ratios (Figure 2b), as increasing β-lactamase can then effectively titrate out the low concentration of clavulanate and restore the resistance phenotype.

### Adaptation of *E. coli* to drug environments

We next tested our theoretical predictions by conducting experimental evolution in 15 drug environments, corresponding to 5 differing amoxicillin proportions (p), each at 3 dose strengths (V) (Figure 2c). Importantly amoxicillin proportion and dose strength (strength of inhibition in the ancestor) varied independently across these 15 environments. After 6 passages (approximately 40 generations) each evolved population was assayed for growth in the drug environment it was selected in. The growth of populations, in their drug environment, was greater for populations selected in high amoxicillin ratios than those selected in low amoxicillin ratios (figure 3a), even though the different drug ratios showed similar efficacy on the ancestral strain (figure S2). The slower adaptation in low amoxicillin lines can also be seen over the course of the whole selection experiment, which ran for 12 passages (figure S4).

**Figure 3:**
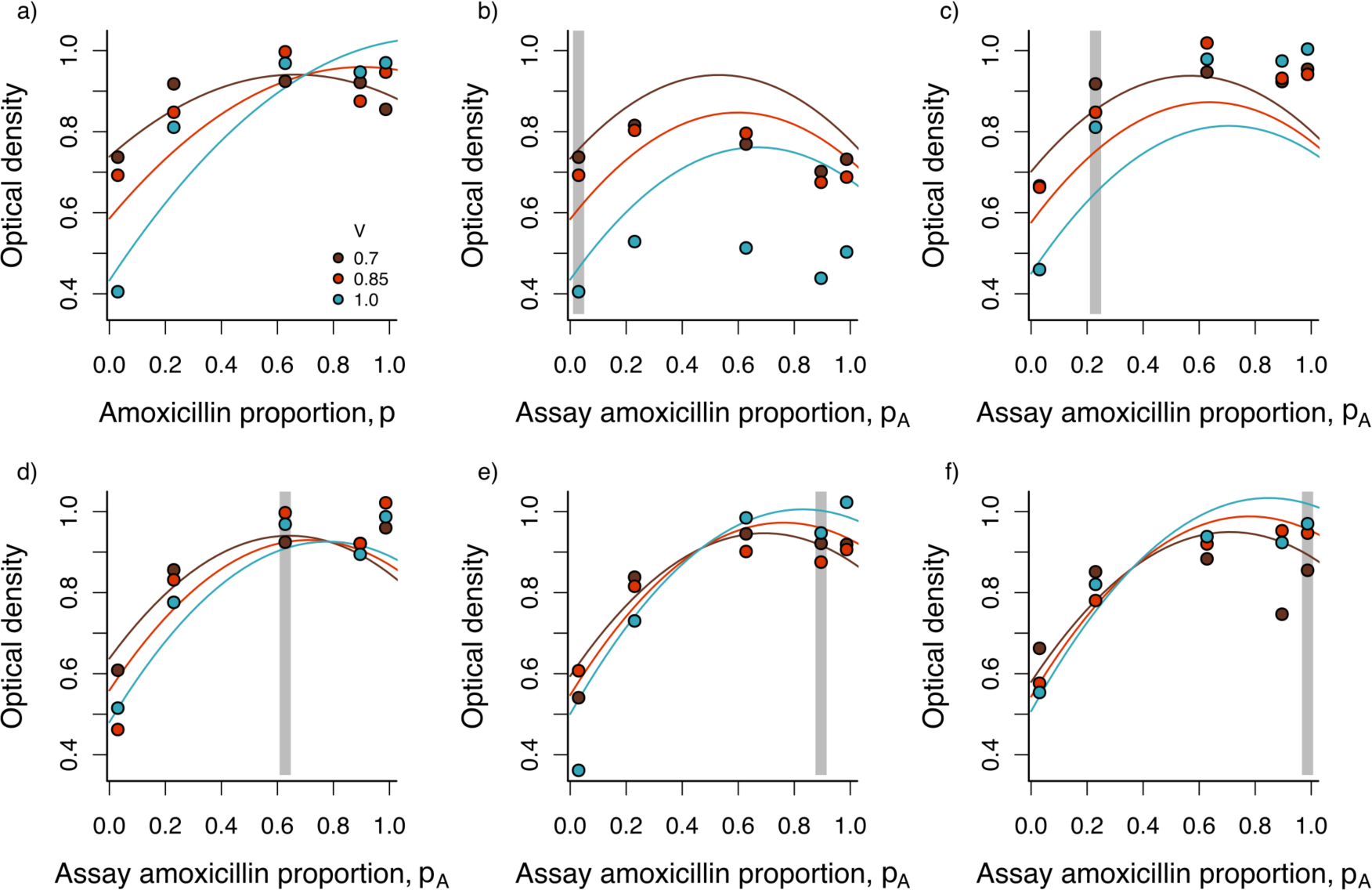
Differences in the adaptation and specificity of resistance for lines evolved under different amoxicillin proportions (*p*). a) shows the growth of each population its selective environment. Panels b-f) show lines evolved at different amoxicillin proportions (p_S_) from a) low *p*_*S*_ to f) high *p*_*S*_ (as depicted in figure 2c). Each line was assayed for growth at alternate drug ratios (*p*_*A*_). Colours differentiate dose strengths for both selection history and assay (*V*_*S*_ = *V*_*A*_). Solid curves show predicted values from the fixed effects of a minimal linear mixed effects model detailed in table S3. The grey regions indicate the drug ratio that lines were selected against, the points in these regions are shown together in panel a). Points are the mean of three replicate evolved lines.

### Cross resistance between drug environments

Next, we explored how adaptation to one drug environment influenced growth across distinct drug environments. The populations that had been selected for 6 passages were exposed to alternate drug environments along the two variables of drug environment; amoxicillin proportion and dose strength.

By varying assay amoxicillin proportion (p_A_) we found that adaptation to a high clavulanate environment (low p_S_, Figure 3b) leads to poor growth across all drug environments. Even the complex statistical model presented here underestimates the extent that lines evolved to high clavulanate treatments have impaired growth. In contrast, adaptation to high amoxicillin environment (high p_S_, Figure 3d-f) leads to performance that is more sensitive to the assay environment p_A_, with high growth in the environment of adaptation and poor growth in a low amoxicillin environment, suggesting higher specificity of resistance in these lines.

By varying the assay dose (V_A_, Figure 4) we found that adaptation to a higher dose strength environment (high Vs, Figure 4c) leads to a reduced sensitivity to increasing assay dose (Figure 4). The lines selected to different drug ratios behave similarly when assay dose strengths are changed. However, strains evolved at high clavulanate proportions do poorly against all dose strengths.

**Figure 4:**
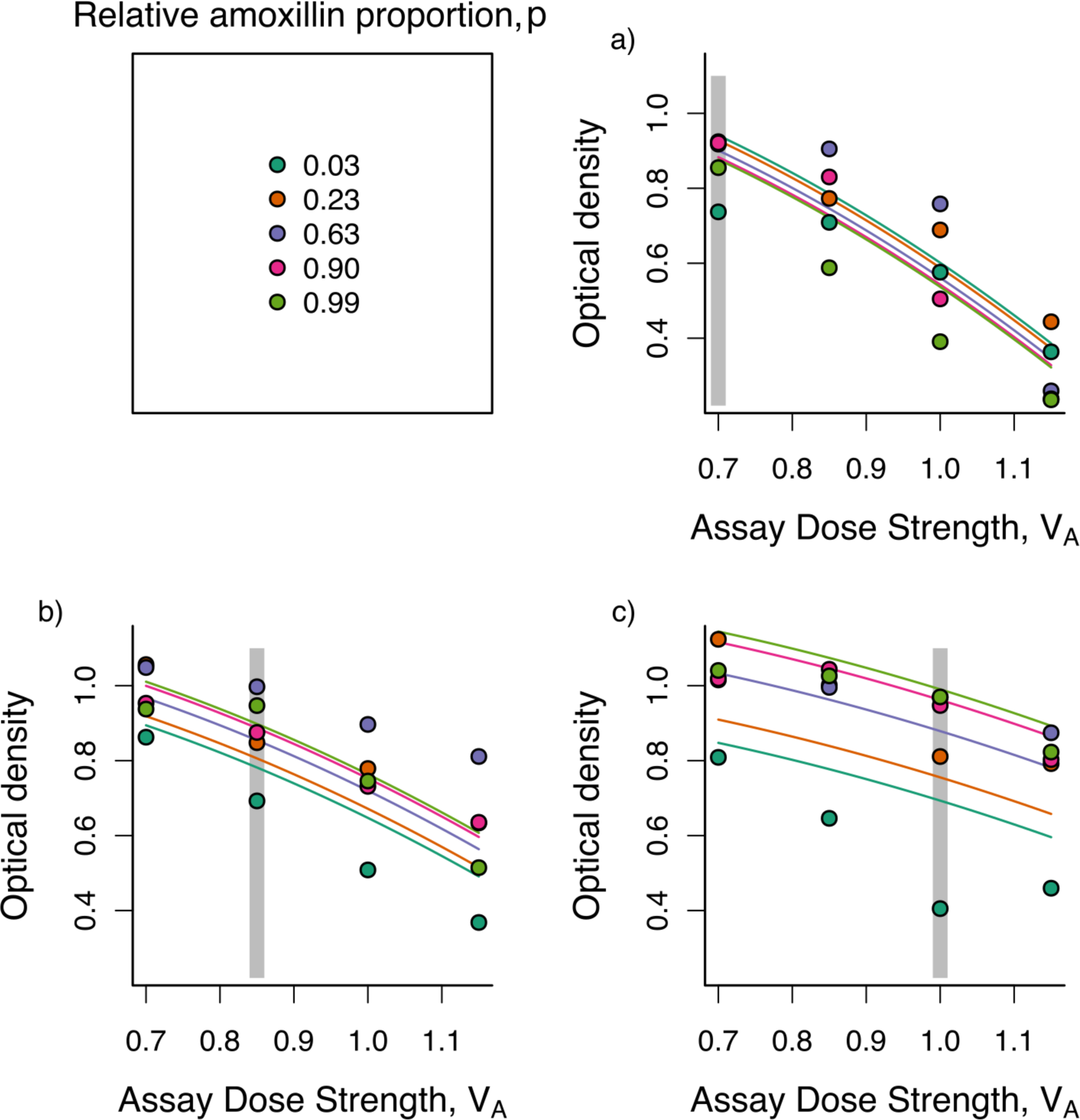
High dose strengths predispose bacteria to growth at even higher drug doses of the same ratio of drugs. Each panel shows lines evolved to dose strengths (*V*_*S*_) of a) 0.7, b) 0.85 and c) 1, assayed for growth at alternate assay dose strengths (*V*_*A*_). Colours differentiate amoxicillin proportion for both selection history and assay (*p*_*S*_ = *p*_*A*_). Solid cures show predicted values from the fixed effects of a minimal linear mixed effects model detailed in Table S4. The grey bars indicate the dose strength (*V*_*S*_) that lines were selected against. Points are the mean of three replicate lines.

### Genetic changes during selection

To cast light on the mechanisms of evolved resistance, we sequenced the 15 populations evolved against high dose strengths of co-amoxiclav. We find a pattern of parallel mutation of the plasmid copy number repression locus *repY* (23), predominantly in the lines evolved against high amoxicillin proportions (Figure 5, Data S1). Mutations affecting porins and efflux pumps, which prevent access of amoxicillin to the cell wall target (24), are also found in multiple lines (Figure 5, Data S1) but are not specifically found in lines evolved to high or low amoxicillin proportions. By using read depth of the plasmid and genome regions to estimate plasmid copy number we find that lines selected against higher amoxicillin proportions evolved higher plasmid copy number (Figure 6, *β* = 2.187, *F*_1,13_ = 20.89, *p*<0.001, robust to removal of outlying point).

**Figure 5:**
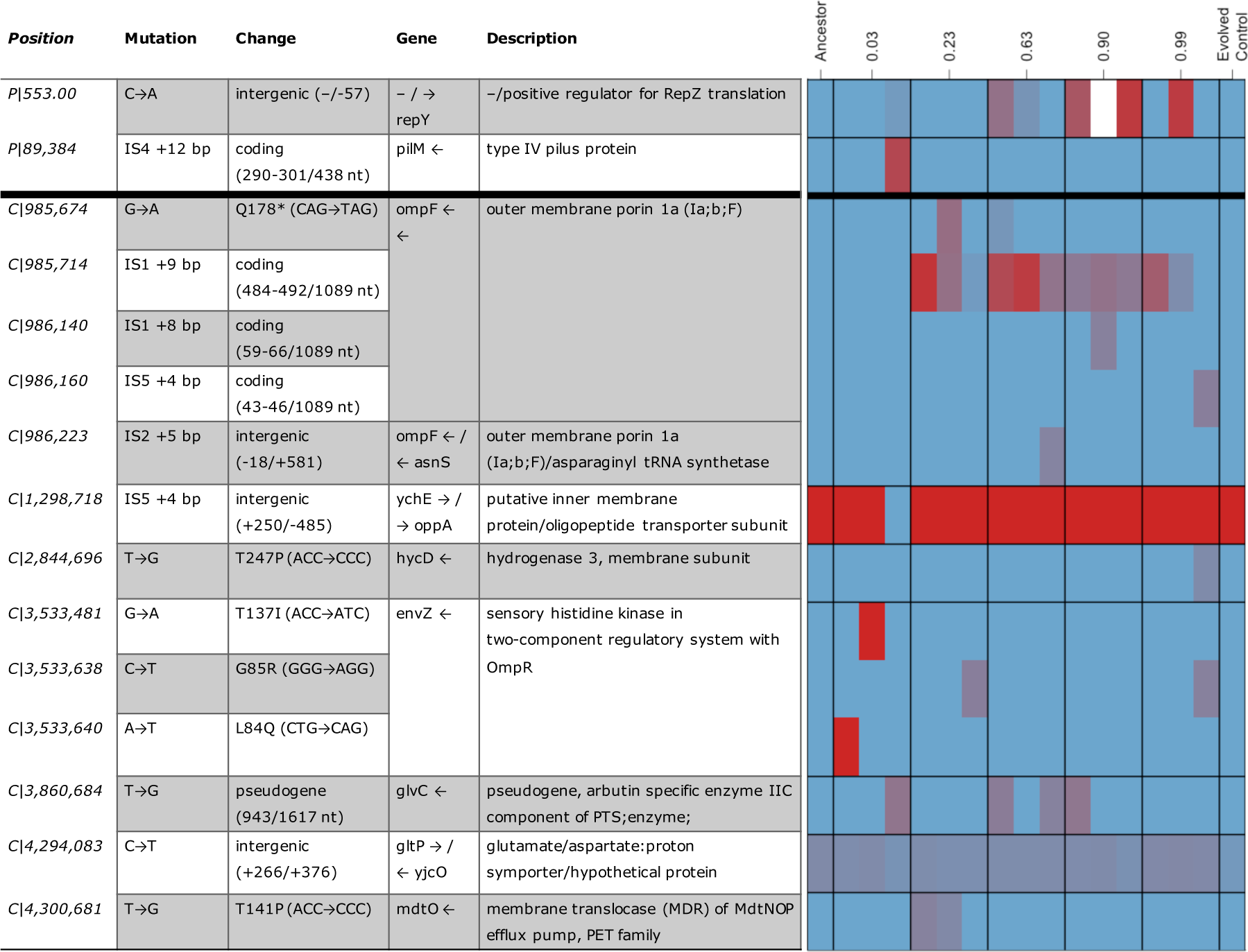
Populations selected for 6 passages have different mechanisms of co-amoxiclav resistance. The table on the left gives detail about the mutations. The frequency of each mutation in all sequenced populations is shown on the left. Red indicates that a mutation is at high frequency and blue indicates that the mutation is at low frequency or absent. The white box indicates that the mutation is present in the population but a frequency is unable to be assigned to it. Mutations are identified by their position and prefixed by P or C for plasmid or chromosome respectively. Only polymorphic mutations that were found at a frequency equal to or greater than 20% in one or more lines are shown. All mutations are listed in the Data S1.

**Figure 6:**
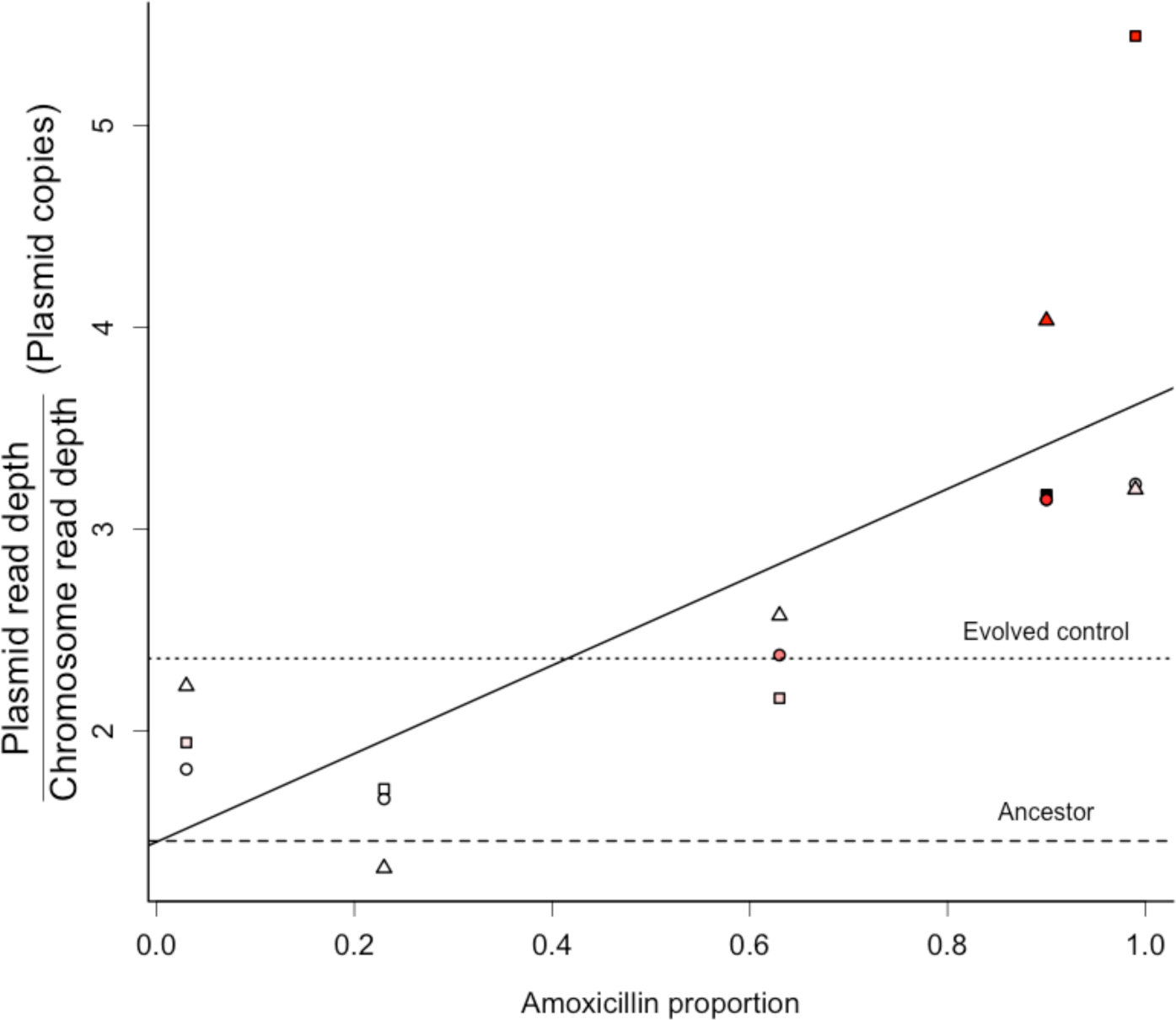
Plasmid copy number is higher in lines selected at high amoxicillin proportions. The copy number of the pCT plasmid in the first second and third replicate populations are denoted by squares, circles and triangles respectively. The strength of the red colour represents the sum of the proportion of all mutations in the *repY* promoter. Replicate 2 of the lines evolved at a relative amoxicillin proportion of 0.90 could not be quantified for the number of mutations at the *repY* promoter and is coloured black.

## Discussion

In this study we have demonstrated that the synergistic interaction between a β-lactam antibiotic and a β-lactamase inhibitor (adjuvant) can lead to distinct phenotypic and genomic paths to resistance evolution in a ratio-dependent manner, with potential consequences for the sustainable management of adjuvant therapies. Recent work has suggested that the ratio of drugs used in combination therapies may affect selection for resistance (25-27). However drug interactions make it difficult to separate the effect of drug ratio from the inhibitory strength of the treatment, which has a well-established effect on the evolution of drug resistance (3). We find that low amoxicillin treatments confer weaker selection for resistance (Figure 3), even when the inhibitory effect of the drug combination on the ancestor is similar. As expected (3), resistance generally evolves faster when the inhibitory effect is greater (figure 3 and 4), however, all lines selected in high clavulanate environments had least resistance, regardless of inhibitory effect (Figure 3a).

Our mathematical model (Figure 2b and S3) suggests that increased β-lactamase expression is more strongly selected with high proportions of amoxicillin, because when clavulanate is not in excess its effect can be titrated out by increasing lactamase expression. On the other hand selection for direct resistance only depends on the inhibitory strength of the drug combination, because this is equivalent to the amoxicillin concentration experienced by the bacterium after some proportion has been broken down by lactamase. Consistent with our model, we found that lines selected at high amoxicillin proportions grew well in high amoxicillin environments but poorly in low amoxicillin environments (Figures 3d-f). These lines had increased plasmid copy number (and thus β-lactamase expression) through parallel mutations in *repY* (Figure 5), which protects against amoxicillin but not in the presence of high levels of clavulanate. On the other hand, lines selected in high clavulanate environments grew poorly, but consistently across all amoxicillin proportions. These lines only acquired direct resistance to amoxicillin through parallel mutations affecting porins and efflux pumps, a resistance mechanism seen across all amoxicillin proportions. This resistance mechanism provides a benefit independent of amoxicillin proportion, as it only depends on amount of non-cleaved amoxicillin. In addition to multiple different mutations in genes with similar functions we find some identical mutations in different lines indicating that specific mutations may be adaptive as is likely the case for *repY* mutations. Mock passaged blank wells showed no evidence of cross contamination. Sequencing of a control evolved line indicated that it was also polymorphic at chromosome position 4,294,083 suggesting this polymorphism was present in the ancestral population or is an artefact of our sequence analysis.

Although the shape of drug interactions have recently been shown to evolve in bacteria (27), to our knowledge this is the first time that this has been reported for antibiotic adjuvant combinations, or that these changes have been linked to the mechanisms of drug action. Our results suggest that dosing regimens with higher amounts of clavulanate will more effectively slow the evolution of resistance by rendering some resistance types ineffective; specifically beta-lactamase dose-response mutations will be less able to titrate out the effect of larger amounts of beta-lactamase inhibitor. Since its introduction the dosage of amoxicillin in co-amoxiclav tablets has increased from 250mg to 875mg, to combat amoxicillin resistance, however the dosage of clavulanate has remained the same at 125mg. There are many other considerations when designing dosing regimens including pharmacokinetics/ pharmacodynamics and toxicity (although amoxicillin and clavulanate are well tolerated, (28, 29), but the potential for resistance is increasingly important. Increased β-lactamase expression is a common resistance mechanism, particularly when plasmid borne (20). Therefore, we suggest that the amount of clavulanate could be increased to reduce selection for increased lactamase expression without affecting the fitness of other resistance mechanisms. This would have the added benefit of reducing selection for plasmid based resistance, which is both easily mobilised and can increase evolvability (20).

As our supply of antibiotics becomes limited there has been greater interest in extending the lifetime of antibiotics through combination with adjuvants, and β-lactams are no exception (9). As antibiotics have been developed for longer than adjuvants we have many β-lactam antibiotics which could be more effective if combined with an adjuvant but relatively few β-lactamase inhibitors to combine them with (15). Therefore, it has been argued that adjuvants should be conserved over antibiotics (13), with the antibiotic component of a combination being replaced when resistance renders it ineffective through direct resistance mechanisms. Our results with co-amoxiclav suggest that using β-lactamase inhibitors at high concentrations would do exactly this by steering resistance away from β-lactamase over-expression and towards direct mechanisms of resistance – at which point the β-lactam component could in principle be replaced. In practice, the potential enrichment of broad-specificity resistance mechanisms will limit the set of replacement options. In general we argue that our ability to manage infections on both the patient and public health scales will require greater investment into the evolutionary consequences of existing and potential treatment regimens.

## Methods

### Strains and media

*Escherichia coli* strain MG1655 containing a naturally occurring pCT plasmid (30) and defective for horizontal transfer due to a mutation in the *trbA* gene (31) was used as the ancestor of all selection lines and is henceforth referred to as the ancestor. The strain was produced in the lab of Dr Ben Raymond (31) and kindly provided. The pCT plasmid is a large naturally occurring plasmid containing the CTX-M-14 extended spectrum β-lactamase. The pCT plasmid is stable, however prior to incubation for experimental evolution and growth dynamics assays the ancestor was grown in the presence of 100µg/ml ampicillin to maintain the pCT plasmid. For phenotyping of experimentally evolved strains pre-culture was without antibiotics to reduce any non-genetic effects of exposure to antibiotic treatment.

Experimental evolution was conducted in a defined minimal medium with the following recipe. M9 medium base (containing 6.78 mg/ml Na_2_HPO_4_, 3 mg/ml KH_2_PO_4_, 0.5 mg/ml NaCl and 1 mg/ml NH_4_Cl) supplemented with 1mM MgSO4, 0.1mM CaCl2, 0.4% (v/v) glycerol, 0.02% casamino acids, 0.5µg/ml thiamine and Hutners trace elements (32) at 1X concentration. Initial checkerboard assays were conducted in Luria Bertani (LB) broth.

Clavulanate (in the form of potassium clavulanate, Fluca analytical) and amoxicillin (LKT laboratories) were supplied in powdered forms, stored at 4°C and used to make stocks in deionised water. These stocks were stored at 4°C according to manufacturer’s instructions, liquid stocks were not kept for longer than 14 days to minimise degradation of the compounds.

To test antibiotic sensitivity of the ancestral strain the ancestor was grown for 22 hours in LB broth in the presence of increasing clavulanate and amoxicillin, at all possible combinations of the two drug concentrations (checkerboard assay, Figures 1b and S1). From these 5 different ratios of amoxicillin and clavulanate, as well as associated iso-inhibitory doses were identified for each ratio. These were tested in minimal medium to confirm that growth was not significantly affected by drug ratio, but was affected by the strength of the drug dose (figure S2). The chosen concentrations and relationships between them are shown in Figure 2c.

### Experimental evolution

To test its ability to adapt to different drug doses, *E coli* was selected against varying drug regimens defined by the relative proportion of amoxicillin (p Selection, p_s_) and dose strength (V_s_), as in figure 2c. A mid exponential culture the ancestor was washed and diluted in minimal medium. This was aliquotted into 48 wells in the centre of a 96 well plate, which were then made up to a final volume of 200µl by adding reconstituted clavulanate and amoxicillin, starting densities were OD = 0.01. Experimental evolution lines were set up corresponding to 5 drug ratios at 3 different strengths, plus one line which was not exposed to drugs, each replicated 3 times (48 independent lines), plus 3 replicate sterile wells with no drugs (which were still passaged). Plates were incubated statically at 37°C for 22 hours for each passage. After each growth cycle wells were mixed using a pipette to re-suspend any clumps of bacteria. The optical density of the wells was then measured and used to transfer cells to a fresh microwell plate so that each line started at an OD (600nm) of 0.01. Experimental evolution was performed for 12 passages (corresponding to approximately 84 generations). Lines were frozen every 3 passages by adding 100µl of a 1:1 LB:glycerol mixture to the remaining culture after the transfer had been performed, these were then frozen at-80°C.

### Measuring cross resistance between drug environments

Although final density was measured at the end of each passage, the growth of frozen samples was used primarily used to assess variation in resistance phenotypes across populations (removing long term physiological effects of antibiotic exposure). To assay evolutionary change in response to drug combinations, for each line the population after 6 passages (chosen because this is when there was most diversity in how lines had adapted to their environment) was revived by overnight growth in LB. Each line of selection was assayed for growth in new drug environments (Assay environments, *p*_*A*_, *V*_*A*_). The differing drug environments that selection lines were assayed against either kept amoxicillin proportion the same (*p*_*A*_ = *p*_*S*_) and varied dose strength (*V*_*A*_) or kept dose strength the same (*V*_*A*_ = *V*_*S*_) and varied amoxicillin proportion (*p*_*S*_). When varying dose strength an increased dose of 1.15 times the maximum dose was also used (*V*_*A*_=1.15). Otherwise all conditions were the same, strains were grown in minimal media statically for 22 hours at 37°C and mixed prior to measuring optical density.

This was a large experiment so selection lines were randomly blocked across the central wells of nine 96 well plates. Each plate had three blank wells and one well containing each of the 3 control lines selected in the absence of drugs and assayed in the absence of drugs. There was small but significant variation in the growth of control lines across plates so OD values were for each plate were corrected using the growth of controls (in the absence of drugs). *p*_*S*_ is undefined for the control lines (selected without drugs) so these were tested against every different drug environment.

### Statistics

All statistics were performed in R (33). Full models were produced using relevant main effects and interactions (statistical tables in supplementary information). Fixed effects models were fitted using the glm function with a Gaussian error distribution and identity link function. For data sets where multiple measures were taken from each strain, a mixed effects model was used to take into account the effect of strain as a random effect, this was fitted using the lme function in the nlme package (34). As there were many explanatory variables for this data, 2 models are fitted to 2 subsets of the data. One data set includes all data where strains are tested for resistance to the same dose strength they are selected against (at varying drug ratios). This will investigate whether resistance evolved to one drug ratio confers resistance to other drug ratios. The other set includes all data where strains are tested for resistance to the same ratio they were selected against (at different dose strengths). This model investigates whether resistance selected at one dose strength confers resistance to others. Both these data sets include the 45 data points (one per evolved line) where both ratio and strength of assay are the same as those for selection (i.e the selection conditions for that line).

The maximal model was reduced to a minimal model in a stepwise manner. At each step of model reduction all terms that were not currently included in an interaction were tested for significance as grounds for including them in a model. Terms were dropped if the result of an F test (for fixed effects models) or likelihood ratio test (for mixed effects models) comparison of models with and without the term of interest was not significant at α=0.05 (i.e. accepting the null hypothesis of no significant effect of the term). At each step only one term could be dropped so where several effects were non-significant the new model with the lowest AiC (Akaike information criterion) was chosen as the best model reduction at that step. When no terms (not included in higher order interactions) could be dropped without a significant effect this was considered the minimal model. Statistical support for all terms in the minimal models was assessed as above but through comparison of the minimal model and the minimal model with the term dropped. Full statistical results are reported in statistical tables in the supplementary information.

### Sequencing and Bioinformatics

To test whether different drug ratios select for different resistance mutations we sequenced evolved populations selected against the highest dose strengths after 6 passages of experimental evolution. The ancestral strain and one of the 3 populations evolved in the absence of drugs was also sequenced. Library preparation and paired end MiSeq sequencing was performed by Edinburgh genomics. Obtained sequences were aligned to both the MG1655 reference (35) and the pCT plasmid reference (30) and polymorphisms identified using breseq in polymorphism mode using default parameters (36).

